# Modular low-cost homemade radiation shields for forest ecology field experiments

**DOI:** 10.1101/2020.08.03.234344

**Authors:** N.S. Morales

## Abstract

1. Abiotic variables are important in understanding several forest ecosystems processes. Hence, temporal, accurate, and precise measurements are vital to any experiment including abiotic variables as part of its objectives.
2. In general, field experiments require several types of sensors to represent intra-site variability. For temperature and humidity measurements in particular, accessories to decrease biases such as radiation shields are indispensable. However, this kind of experimental setup can make project costs escalate quickly.
3. Low-cost alternatives to commercially available radiation shields have proven to be adequate in their ability to decrease measurement bias. Despite this advantage (low cost), the materials used in some of these cannot last long under field conditions (e.g. cardboard, aluminium foil).
4. A custom-made low-cost solution was designed to withstand rugged conditions to perform temporal measurements using sensors. The radiation shields proved to meet the technical and budgetary needs of the project while lasting the entire duration of the project (approx. 3 years). At the end of the project they were still in good enough condition to use in future studies, and they were donated to the School of Environmental Sciences (University of Auckland).

## Introduction

The importance of abiotic variables on forest processes require accurate and precise temporal measurements. Sensors and dataloggers are a crucial part of most forest ecology studies, as changes in abiotic conditions can have a huge impact on different forest ecosystem processes such as plant regeneration, biomass decomposition rates, flammability of biomass, etc (e.g. Ewers & Banks-Leite 2013, Rossa 2017). In addition to the sensors, there are other accessories that need to be considered to minimize measurement errors. For example, the potential bias on temperature and humidity measurements caused by solar radiation striking the sensors directly, makes the use of proper shielding critical. To avoid biased measurements, sensors are usually placed inside a structure commonly referred as a radiation shield (Hubbard, Lin & Walter-Shea 2001). Radiation shields come in a wide variety of shapes and forms, but their main functions are to shield against direct sunlight while allowing appropriate ventilation for accurate sensor reading.

The cost of sensors plus radiation shields can transform an experimental design into a prohibitive expensive endeavour (Terando, Prado & Youngsteadt 2018). In general, a field experiment will require several of these instruments to obtain a representative data trend from each site. Although there are radiation shield designs that can be easily reproduced and that there are comparable, performance-wise, to commercially available ones, the materials used are not sturdy enough to endure extended periods in the field (e.g. cardboard, aluminium foil) (Tarara & Hoheisel 2007). In addition, both commercial radiation shields and the currently available alternatives require the researcher to carry several, often bulky, structures to replace potentially faulty shields and/or sensors for the duration of the study. Currently, there are no alternatives that allow researchers to carry only spare parts. The lack of practical alternatives can pressure researchers to reduce sample sizes to adjust for minimal research funding, sacrificing appropriate study confidence levels. This challenge is difficult to resolve, especially by researchers in low-income countries, where there is a necessity to decrease the cost of the equipment needed and at the same time maintain the statistical soundness of the experiments.

The methodology described here addresses the challenges presented above. I designed and built a new low-cost modular gill radiation shield using sturdy materials that can fulfil the requirements of forest ecology experiments in the field. The proposed solution was used in a restoration and forest fragmentation field experiment context. A total number of 27 of these radiation shields together with temperature and humidity sensors (iButtons®) were used in the field for approximately three years (see Morales, Perry & Burns 2016).

## Description and implementation

### Radiation shield design

The radiation shield design was conceived to be modular and robust. The final design was easy to dismantle, meaning that each individual part could be removed and replaced, allowing repairs in the in the field if necessary. The gill radiation shield design consisted of set of concentric plastic plates fixed together by bolts (Fig. 1). The custom design proposed here was devised for iButtons® sensors. However, the methodology described here is generic and flexible enough that with small changes to the dimensions of the required materials, one could hypothetically fit a sensor of any size.

**Figure 1.**
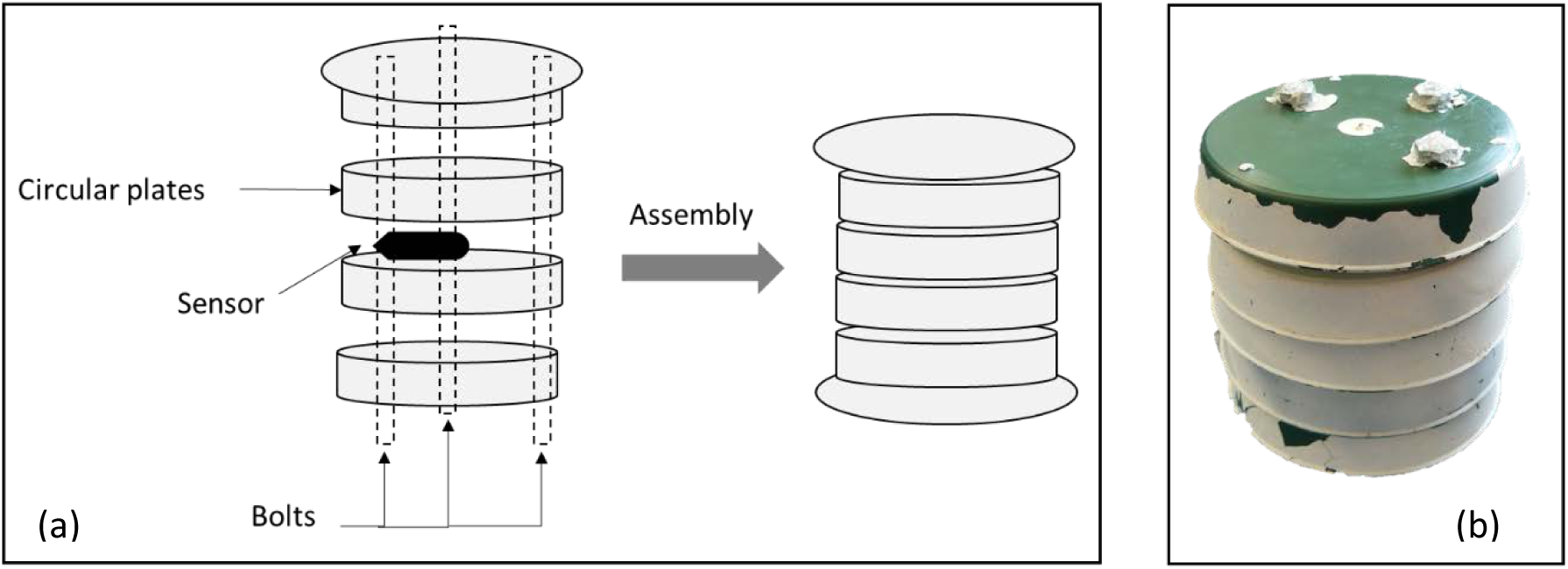
Diagram showing the radiation shield design: (a) main parts used in the assembly and a photo of the final product after three years of use in the field. In this case the plastic plate used were originally green and were painted white hence the observed paint peeling.

### Materials

Materials used in the assembly of these radiation shields are presented in Figure 2. The selection of materials was based on three main criteria: 1) widely available, 2) inexpensive and 3) reusable. One of the most important components are the plastic saucers. I chose a type of plastic that was flexible but not brittle (10 cm diameter, 18 mm height; preferably white). I only managed to find green saucers, but the company was open to make them in white if needed. Three bolts with nuts were used per radiation shield (size can vary depending on the how many plates you need to add). Galvanized bolts were used, and no rust was observed after three years in the field (temperate type forest with high average rainfall). However, if you were to use them in a warmer and more humid type of climate, stainless steel would likely be more adequate.

**Figure 2.**
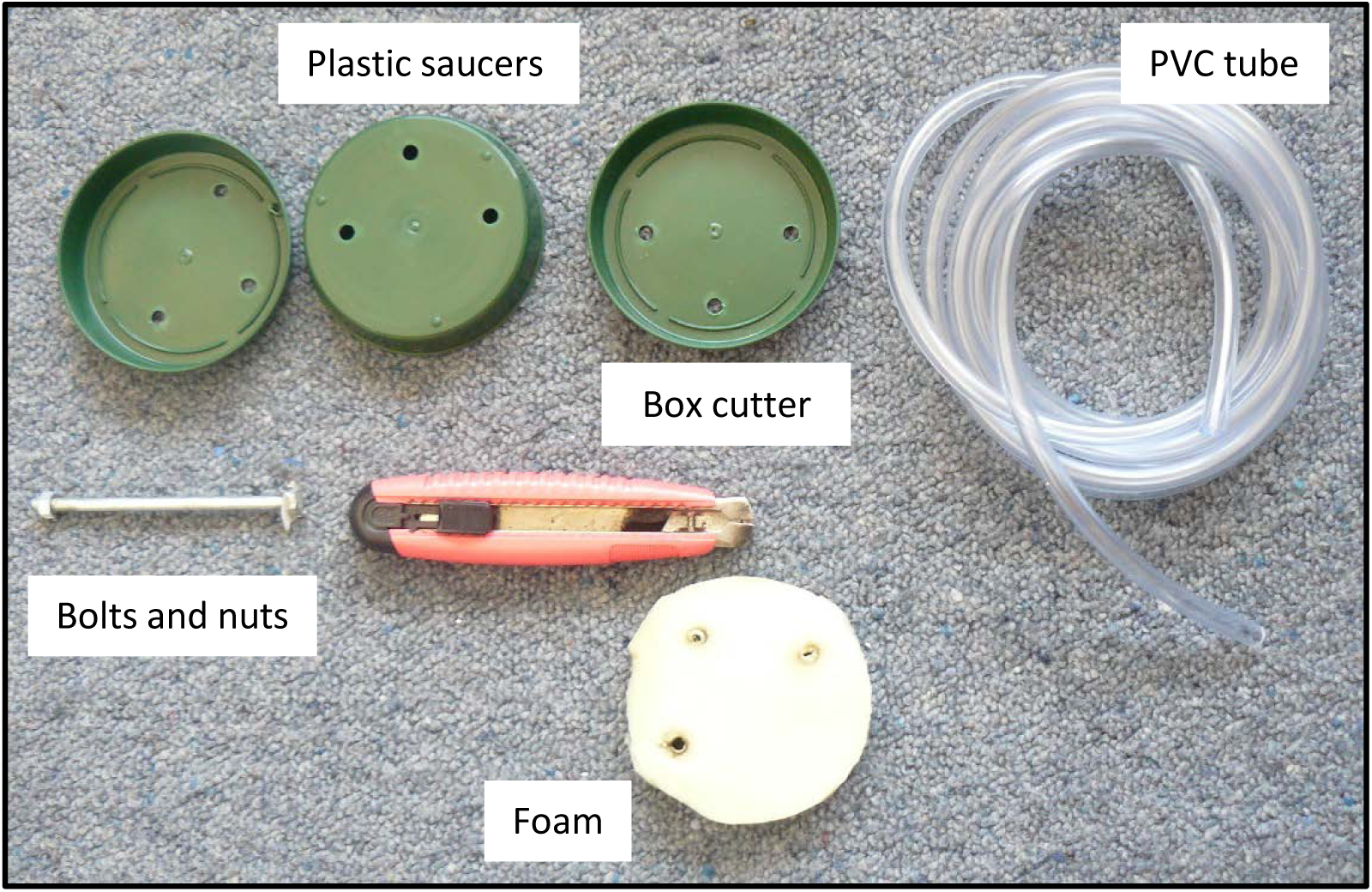
Materials necessary to build the radiation shields

A second important component is flexible PVC tube (the type generally used in aquariums), that is of the same diameter as your bolts. Also, some foam is needed to insulate the top part/saucer of the radiation shield. There is no specific type of foam to use, even you can recycle packaging foam to this purpose. To avoid water entering through the bolts you can use silicon sealant or hot glue. The approximate cost of each radiation shield is $5 USD, but this cost can vary depending on your country of residence, changes in the quality of materials or including recycling of disposable materials (e.g. foam). Nonetheless, when compared with a commercial radiation shield, where prices can range from $20 - $150 USD or more, the alternative presented here will likely be much more cost effective.

### Preparation of plastic saucers

For a radiation shield large enough to fit the sensor while allowing air flow, you will need approximately five plastic saucers. The number can be adjusted depending on your needs, but I do not recommend using less than five saucers. You will need to make three holes in each of the plastic saucers. The most important part of this process is the alignment of the holes. If holes are not properly aligned you will not be able to assemble the radiation shield. To ensure the holes are in the same position in each of the plates, prepare a template, mark the holes and make the holes in one saucer first. Once you are certain of where the holes will be, the easiest way it is to group the plastic saucers in a bunch of five and use a handheld drill or a drill press to make the holes in them all at once. The use of a drill is optional but recommended. If you do not have one, ask your faculty or school workshop if they have one you can borrow. When all the holes are done, you will need to cut a circle out in the centre of three of the plastic saucers, while 2 remain intact (Fig. 3). You will obtain better results if you heat the cutter first. Finally, if your saucers are not white paint the plastic saucers with white spray paint (Fig. 3). Purchasing white saucers is preferred because depending on the material of the saucers and type of paint, the paint will start to peel off in approximately six months (depending on local conditions). If paint starts to peel of, you will need to repeat the painting process. This process, however, is inexpensive and simple to carry out.

**Figure 3.**
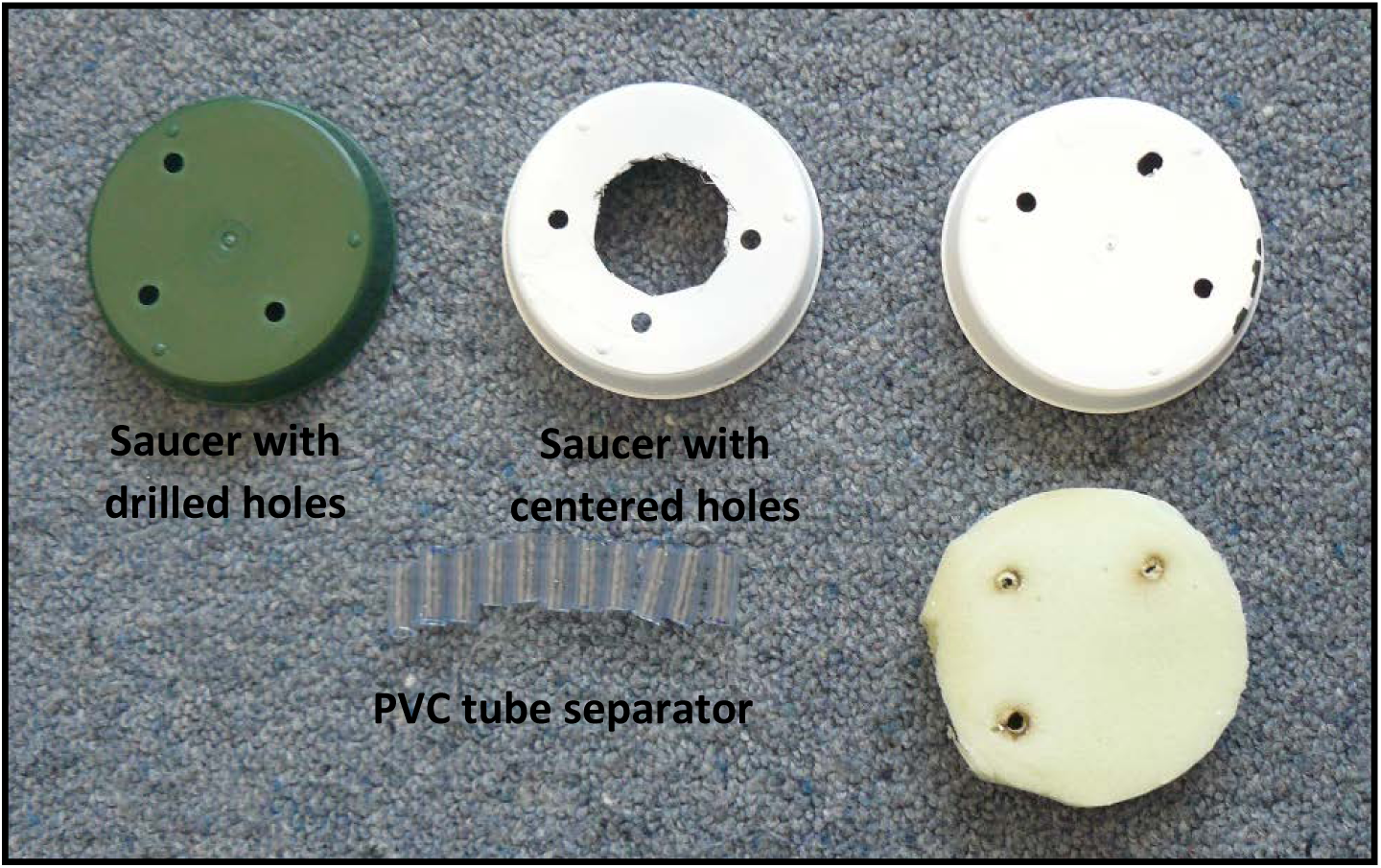
Saucer with drill holes and the two different settings of saucer use to complete a radiation shield.

### Adding insulation

You will need to add a layer of insulation to the radiation shield. In this model, the insulation will for inside the “top” saucer to decrease the effect of direct light which can have a negative on the measurements (Nakamura & Mahrt 2005). First, you will need to cut out circular shapes from the soft foam of the same diameter as the plastic saucers. You will need only one of these foam circles per radiation shield. Take one of the “complete” plastic saucers (not the ones with a circle in the middle) and place the foam in it. You can glue the foam to the saucer, but it is not compulsory (Fig. 4).

**Figure 4.**
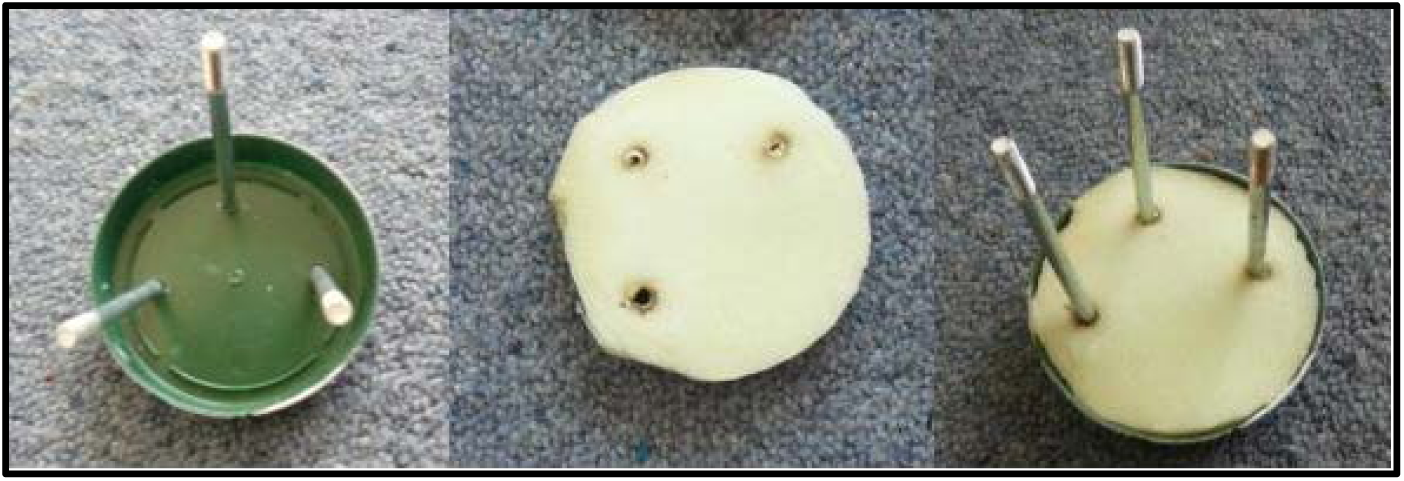
Proper fit of foam circle in the plastic saucer

### Radiation shield assembly

Push the bolts through the opening of the first saucer (with no hole in the centre) and push your circular piece of foam to fit as seen in Fig. 4. To ensure adequate ventilation, there must be a standard separation between each saucer that makes up the radiation shield (Nakamura & Mahrt 2005). To make separators, cut the PVC tube with a hot box cutter into small sections (approx. 2 cm in length, but length will depend on the size of your radiation shield) (Fig. 3). Slide a separator onto each bolt and push it down close to the foam. Next, place one of the plastic saucers with a circle in the middle through the small holes for the bolts and put another set of “separators” on each bolt. I usually put the sensor in at this level using a fob specially designed to hold an iButtons®. Repeat the process as needed to complete the radiation shield. To complete your radiation shield you should use the second plastic saucer with no hole to finish the structure. Finally, tighten the nuts at the end of each bolt (Fig. 5).

**Figure 5.**
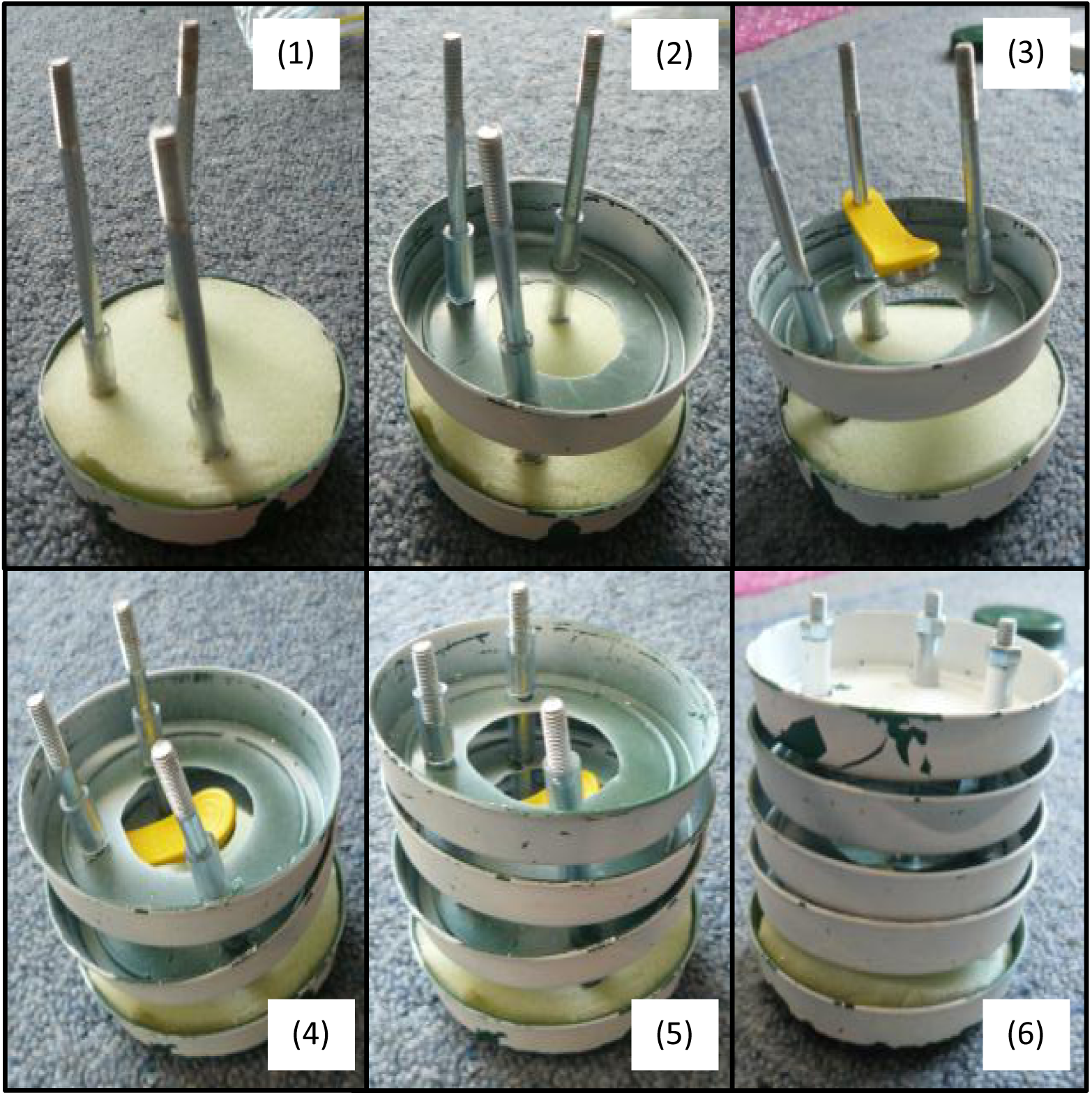
Photographic sequence showing the assembly of the different plastic saucers and how to fit the iButtons® in the structure (sequence (1)-(6))

### Radiation shield testing

I compared the performance of the handmade radiation shield with a commercially available one (same gill design), during spring for a period of 24 hours exposed to the elements, including direct sunlight. I did not find significant differences on temperature between radiation shields; *t*(8)= -1.51, *p*= 0.169 (Fig. 6).

**Figure 6.**
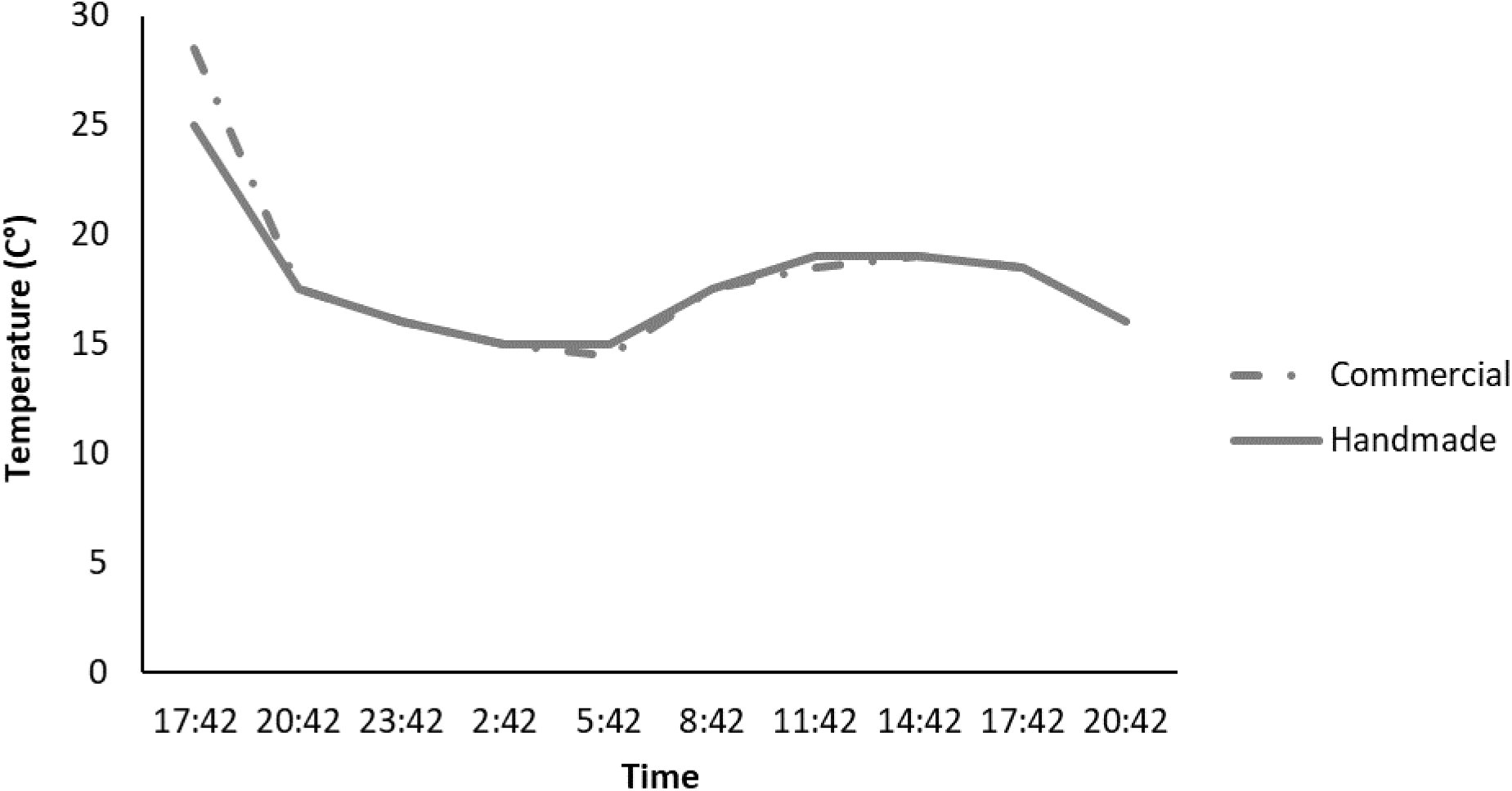
Graph comparing temperature (C°) and measurements during a period of 24 hours for the tested radiation shields. The first measurement was not including in the analysis because it was made while I was setting up the radiation shields.

### Field setup

Once you have finished assembling the radiation shield, you will need to seal it to prevent water and other debris from entering the housing. Using silicone or hot glue, make sure to seal around the “heads” of the bolts on the top of the radiation shield. Now they are ready to be put in the field. You can fix the radiation shield to a pole with cable ties which are highly durable in the field, using the bolts for this purpose (Fig. 7). Make sure to leave space for air flow so do not fix them too tight. Attaching the radiation shields to large trees, for example, is not recommended. Not only does this limit airflow and can therefore bias results, it also exposes radiation shields and sensors to increased humidity from the tree, and vegetation debris. The radiation shields described here were used in a long-term field study and were left in the field for almost 3 years. During this time no maintenance was required, except the occasional touch up of white paint. The sensors were not only exposed to the elements but also to large mammals (e.g. cows). In one such instance the pole where the radiation shield was attached was knocked over, the radiation shield showed visible signs of being kicked/trampled on. However, there was minimal damage and I was able to continue using the radiation shield for the duration of the study. The design proposed here allows these radiation shields to perform quite well under wet conditions.

**Figure 7.**
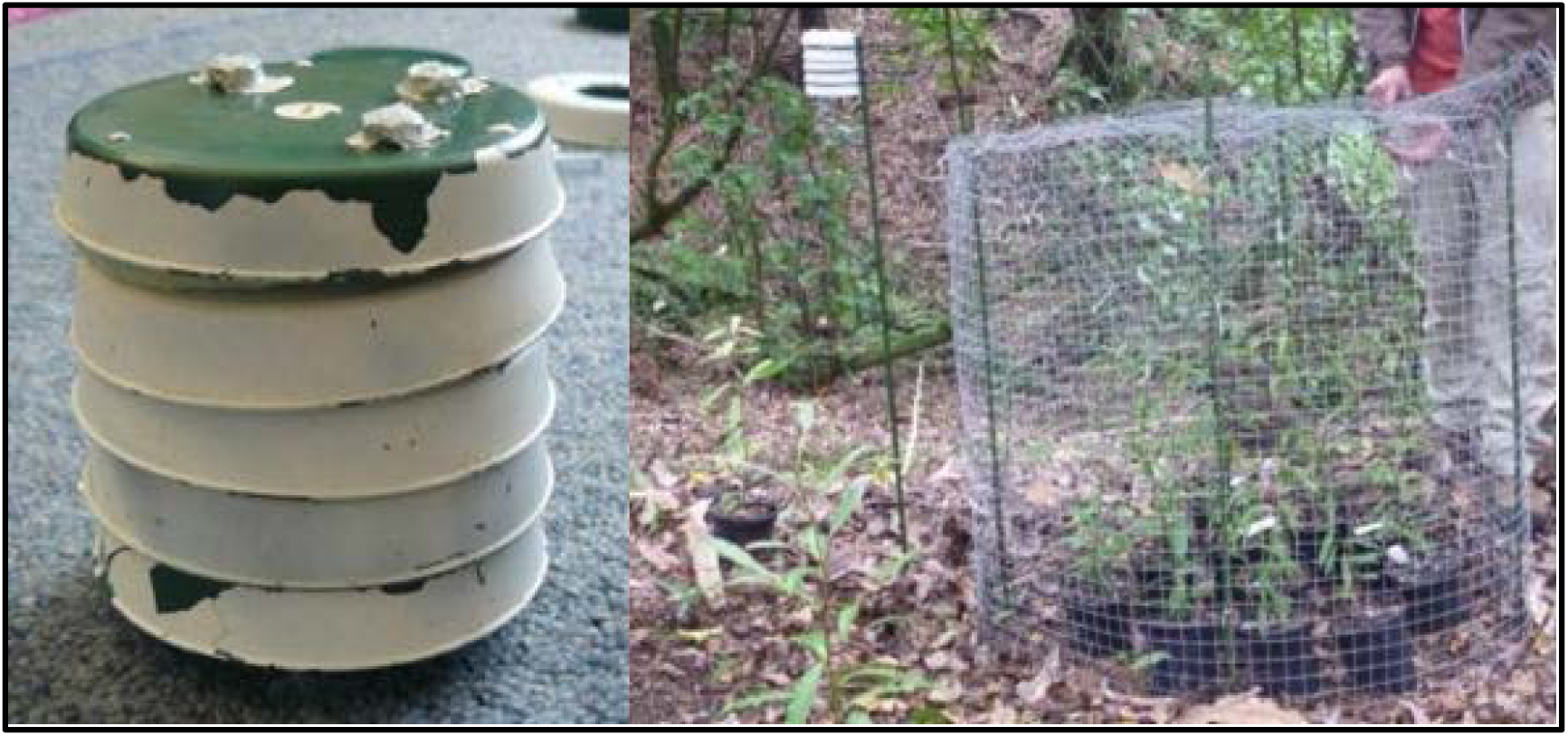
Radiation shield finished and installed in the field

## Conclusion

The custom-made radiation shields presented here are a practical and feasible alternative to the commercially available kind. They proved to be dependable and sturdy enough to withstand a wide range of climatic conditions and trampling by large mammals. Thanks to the modular design, it was only necessary to carry the elements necessary to perform repairs, avoiding the need to carry complete spare radiation shields on each trip to the field. One downfall of the design was the colour of the plastic saucers, I would recommend procuring white saucers instead of painting them. Paint needed to be retouched in some shields during the experiment which is a mayor point to considerer if you are not visiting your experimental sites often. Overall, this design worked under field conditions, meeting technical expectations and the project’s limited budget.

## Acknowledgments

I would like to thank George Perry and Brendan Hall for their support during the design and application of the solution proposed here. I would also like to thank the School of Environment at University of Auckland and Sandra Anderson for her kind encouragement to write this manuscript. Also, I would like to thank Giselle Muschett for helping in polishing this manuscript.

